# CardioPy: An open-source heart rate variability toolkit for single-lead EKG

**DOI:** 10.1101/2020.10.06.328856

**Authors:** Jackie L. Gottshall, Natasha Recoder, Nicholas D. Schiff

## Abstract

**Background and Objective:** Heart rate variability (HRV) is a promising clinical marker of health and disease. Although HRV methodology is relatively straightforward, accurate detection of R-peaks remains a significant methodological challenge; this is especially true for single-lead EKG signals, which are routinely collected alongside EEG monitoring and for which few software options exist. Most developed algorithms with favorable R-peak detection profiles require significant mathematical and computational proficiency for implementation, providing a significant barrier for clinical research. Our objective was to address these challenges by developing a simple, free, and open-source software package for HRV analysis of single-lead EKG signals.

**Methods:** CardioPy was developed in python and optimized for short-term (5-minute) single-lead EKG recordings. CardioPy’s R-peak detection trades full automation and algorithmic complexity for an adaptive thresholding mechanism, manual artifact removal and parameter adjustment. Standard time and frequency domain analyses are included, such that CardioPy may be used as a stand-alone HRV analysis package. An example use-case of HRV across wakefulness and sleep is presented and results validated against the widely used Kubios HRV software.

**Results:** HRV analyses were conducted in 66 EKG segments collected from five healthy individuals. Parameter optimization was conducted or each segment, requiring ~1-3 minutes of manual inspection time. With optimization, CardioPy’s R-peak detection algorithm achieved a mean sensitivity of 100.0% (SD 0.05%) and positive predictive value of 99.8% (SD 0.20%). HRV results closely matched those produced by Kubios HRV, both by eye and by quantitative comparison; CardioPy power spectra explained an average of 99.7% (SD 0.50%) of the variance present in Kubios spectra. HRV analyses showed significant group differences between brain states; SDNN, low frequency power, and low frequency-to-high frequency ratio were reduced in slow wave sleep compared to wakefulness.

**Conclusions:** CardioPy provides an accessible and transparent tool for HRV analyses. Manual parameter optimization and artifact removal allow granular control over data quality and a highly reproducible analytic pipeline, despite additional time requirements. Future versions are slated to include automatic parameter optimization and a graphical user interface, further reducing analysis time and improving accessibility.

## 1. INTRODUCTION

Heart rate variability (HRV) is a measure of fluctuations in cardiac beat-to-beat interval lengths produced by heart-brain interactions that regulate the autonomic nervous system [1]. As such, it provides a functional index of nervous system regulation across a diversity of applications ranging from athletic fitness [2] to psychological stress [3] to cardiac mortality risk [4]. Single and multi-lead electrocardiograph (ECG/EKG) signals required for HRV analysis are routinely collected during many clinical applications, providing a wealth of raw data available for clinical research.

A number of HRV software programs have been developed and scientifically validated through the peer-review process (**Table 1**) [5–9]. Each program outlined in **Table 1** calculates various time and frequency domain analyses standard to HRV estimations. Each of the above-described metrics is calculated from the base tachogram of NN intervals, derived as the differences between subsequent R-peaks in the EKG QRS complex after removal (or “cleaning”) of ectopic and abnormal beats. Time domain analyses (**Table 2**) are calculated directly from the tachogram of interbeat intervals; these metrics quantify the overall variability of intervals between successive heart beats, while frequency metrics allow for characterization of the origins of this variability [10]. Frequency domain analyses (**Table 2)** are calculated from spectral density of the interbeat interval tachogram in discrete frequency bands, standardized into ultra-low-frequency (ULF, ≤0.003 Hz), very-low-frequency (VLF, 0.003-0.04 Hz), low-frequency (LF, 0.04-0.15 Hz), and high-frequency (HF, 0.15-0.40 Hz). The ratio between LF and HF components has been proposed to represent the interplay between sympathetic and parasympathetic nervous system activation, with sympathetic tone tending toward the low-frequency component and parasympathetic tone toward the high-frequency component [11]. Additionally, a number of tools calculate non-linear analyses, which approximate the statistical complexity of the NN tachogram.

**Table 1.** Overview of commonly used peer-reviewed HRV software tools. CardioPy is included in grey shading for a comparative overview.

|  | **Kubios**<br>**HRV**<br>**(Standard)** | **Kubios**<br>**HRV**<br>**(Premium)** | **gHRV** | **HRVanalysis** | **ARTiiFACT** | **PhysioLab** | **CardioPy** |
| --- | --- | --- | --- | --- | --- | --- | --- |
| **Price** | Free | Paid | Free | Free | Free | Free* | Free |
| **Open**<br>**Source** | No | No | Yes | No | No | Yes | Yes |
| **Operating**<br>**System** | Windows,<br>MacOSX,<br>Linux | Windows,<br>MacOSX,<br>Linux | Windows,<br>MacOSX,<br>Linux | Windows | Windows | Windows,<br>MacOSX | Windows,<br>MacOSX,<br>Linux |
| **Language** | MATLAB | MATLAB | Python | MATLAB | MATLAB | MATLAB | Python |
| **Interface** | GUI | GUI | GUI and<br>spreadsheet-<br>like | GUI | GUI | GUI | Command<br>line,<br>Jupyter |
| **R-peak**<br>**Detection** | No | Yes | No | Yes | Yes | No | Yes |
| **HRV**<br>**Analyses** | Time &<br>frequency<br>domain,<br>non-linear | Time &<br>frequency<br>domain,<br>non-linear | Time &<br>frequency<br>domain,<br>non-linear | Time &<br>frequency<br>domain,<br>non-linear | Time &<br>frequency<br>domain | Time &<br>frequency<br>domain | Time &<br>frequency<br>domain |

**Table 2.** Standard Time and Frequency Domain HRV Analyses. Calculations are appropriate for short-term (5 minute) recordings and exclude statistics that require longer data segments. In the CardioPy toolkit, total power is calculated across the spectrum and for each frequency band. Log power, peak power, and % power are calculated for each frequency band. Power (nu) is calculated for LF and HF bands.

| Analysis | Description |
| --- | --- |
| **Time Domain** | Calculated from the cleaned NN tachogram |
| Average interbeat interval | Average of NN intervals (milliseconds) |
| Average Heart Rate | Mean interbeat interval in beats per minute (bpm) |
| Max/Min Heart Rate | Max/min of the sequence of mean heart rates (bpm) calculated over a moving window of 5 NN intervals |
| SDNN | Standard deviation of NN intervals |
| RMSSD | Root mean square of the difference between successive NN intervals |
| pNN20 | Sample percentage of consecutive NN intervals differing by more than 20 milliseconds |
| pNN50 | Sample percentage of consecutive NN intervals differing by more than 50 milliseconds |
| **Frequency Domain** | Calculated from power spectrum of the cleaned, interpolated, and resampled NN tachogram |
| Total power | Total power (milliseconds<sup>2</sup>) |
| Log Power | Log of power |
| Peak Power | Frequency with maximum power (Hz) |
| % Power | Proportion of power across the spectrum |
| Power (nu) | Power expressed as the proportion of combined power in the LF and HF bands (normalized units, nu) |
| LF/HF Ratio | Power in the LF band divided by power in the HF band |

Notably, most validated tools are built in the proprietary MATLAB environment, which may limit transparency and analytic flexibility. Moreover, many require R-peak detection and cleaning to be completed prior to data loading. This provides a particular challenge for analysis of single-lead EKG signals, which are not typically supported by HRV software. To address this gap, we developed CardioPy as a novel R-peak detection and HRV analysis program for single-lead EKG. It is written in python and leverages the vetted functionalities of NumPy, Pandas, and SciPy, allowing for quick and flexible time and frequency domain analyses, as well as a highly reproducible analytic pipeline for production of high-quality scientific studies.

In the following sections we provide an introduction to the CardioPy toolbox. We then apply CardioPy to an example analysis of HRV in five healthy individuals, first validating our results against the widely-used Kubios HRV software, and then examining differences in HRV between wakefulness and sleep states.

## 2. METHODS

### 2.1 Project Vision

CardioPy was born out of the desire to conduct HRV analyses on single-lead EKG data collected from clinical EEG monitoring. Despite significant interest in HRV as a clinically useful physiological metric, we found the practice of conducting HRV analyses on clinical data to require a high bar of domain-specific expertise. The two major obstacles that we encountered were: (1) a scarcity of freely available R-peak detection and cleaning tools and (2) a failure of highly successful published R-peak detection algorithms to supply easy-to-implement code for reproducible analyses. Accordingly, our goal was to produce an HRV analysis toolkit for single-lead EKG that was freely available, open source, and easy to understand and implement.

### 2.2 Usability and Computational Efficiency

CardioPy is written in python, leveraging the language’s readability and wealth of open-source libraries and troubleshooting resources. It requires a python 3 distribution including standard data processing and visualization dependencies, all of which can be obtained through Anaconda (Anaconda Software Distribution). Analyses can be run through any python interpreter, although jupyter notebook use is recommended for analytic documentation and reproducibility.

For structure, CardioPy utilizes a single EKG class allowing for self-contained methods for R-peak detection, analysis, and visualization in matplotlib. Data is stored largely in Pandas DataFrames for easy inspection. The strategic choice of using Pandas data structures in combination with matplotlib visualizations results in an EKG object that is highly readable in exchange for moderate computational efficiency. For this reason, CardioPy is optimized for datasets of approximately 5 minutes in length — the gold standard for short-term EKG recordings (11). While CardioPy may be used for longer datasets, it will likely suffer lagging visualizations and statistical analyses, particularly in the frequency domain.

### 2.3 Data Compatibility & Documentation

Plain text (.txt) data derived from single-lead EKG, either as a single column or as part of a larger EEG file are compatible with CardioPy. Input assumes two rows with column headers specifying both channel and column type, such that line 1 of the EKG column is labeled “EKG” and line 2 “Raw”. CardioPy is compatible with previously cleaned data, allowing for the presence of missing rows and NaN values. Full documentation is available on GitHub (github.com/CardioPy/CardioPy), complete with installation instructions and example analyses.

## 3. RESULTS

The following section provides an overview of CardioPy. First, we explain various CardioPy functions, including R-peak detection and parameter optimization, artifact rejection, HRV analysis, and data export. For parameter optimization, we report algorithm performance in terms of detection sensitivity (percentage of true peaks detected) and positive predictive value (PPV) (percentage of detections that are true peaks). We then provide an example use-case complete with code snippets (**Supplementary Material**) to demonstrate how to implement various functionalities.

### 3.1 R-peak Detection

#### 3.1.1 Detection Algorithm

To detect R-peaks, a dynamic threshold is calculated as the average signal over a moving window (default: 100 milliseconds) shifted upwards by a percentage of the signal value (default: 3.5%). R-peaks are identified as the local maxima between positive- and negative-slope threshold crossings. **Figure 1** demonstrates algorithm performance on a typical EKG signal with default threshold parameters. For this signal, the thresholding algorithm correctly detected 328 out of 328 R-peaks with 0 false detections, resulting in a sensitivity of 100% and a positive predictive value of 100%.

**Figure 1.**
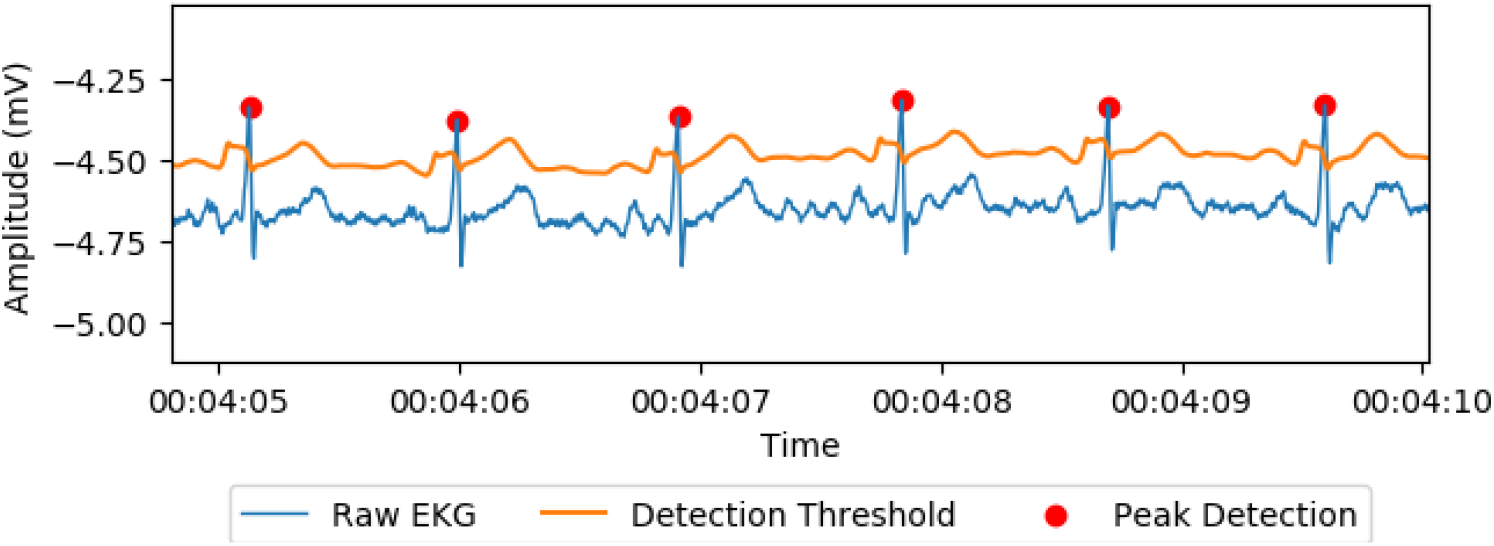
Illustration of R-peak detections. Dynamic threshold and resulting R-peak detections on a standard single-lead EKG dataset. Raw EKG is shown in blue, detection threshold in orange, and peak detections in red.

#### 3.1.2 Algorithm Flexibility

The two main parameters of CardioPy’s detection threshold are easily modified for varying signal amplitude and noise levels. The moving window length can be decreased or increased to produce more or less responsiveness to signal shifts, respectively. Percentage upshift adjustment allows for adaptation to varying R-peak amplitudes between signals. Parameters can be adjusted individually or in tandem. **Figure 2** demonstrates the effects of parameter adjustment on a signal for which the applied parameters performed poorly (sensitivity = 100%, PPV = 60.5%) (**Figure 2, Top Panel)**. In this example, shrinking the moving window size from 300 to 100 milliseconds drastically improved algorithm performance (sensitivity = 100%, PPV = 99.7%) (**Figure 2, Center Panel)**. Alternately, increasing the percentage upshift from 2.0% to 3.5% achieved threshold optimization (sensitivity = 100%, PPV = 100%) **(Figure 2, Bottom Panel)**.

**Figure 2.**
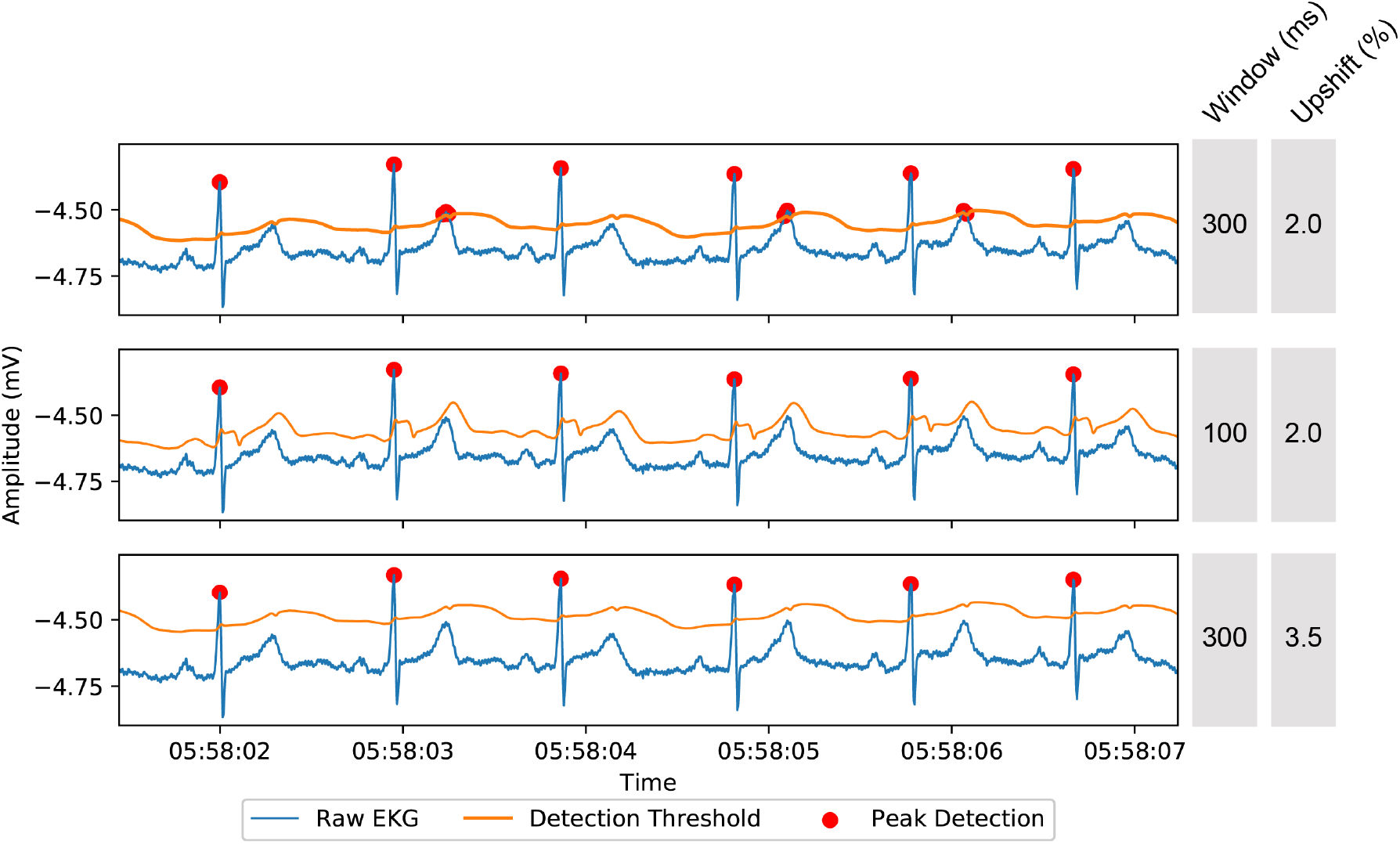
Illustration of threshold parameter modifications. **Top panel)** Raw EKG and detection threshold tracings for a signal with many false peak detections**. Center panel)** Raw signal with moving window size decreased from 300 to 100 milliseconds, resulting increased threshold responsiveness and reduced false peak detections. **Bottom panel)** Raw signal with threshold upshift increased to 3.5% and moving window size set to 300ms, achieving detection optimization. Raw EKG signal is shown in blue, detection threshold in orange, and peak detections in red.

Especially noisy signals can be smoothed with a centered moving average window prior to threshold calculation. **Figure 3** demonstrates an example of smoothing applied to a noisy signal. In this example, the applied parameters often failed to differentiate between signal and noise, resulting in 100% sensitivity and 64.0% PPV. Applying pre-threshold smoothing with a 20-millisecond window optimized detection performance to 100% sensitivity and 100% PPV. As with threshold parameters, length of the smoothing window can be easily modified to accommodate the characteristics of a given dataset.

**Figure 3.**
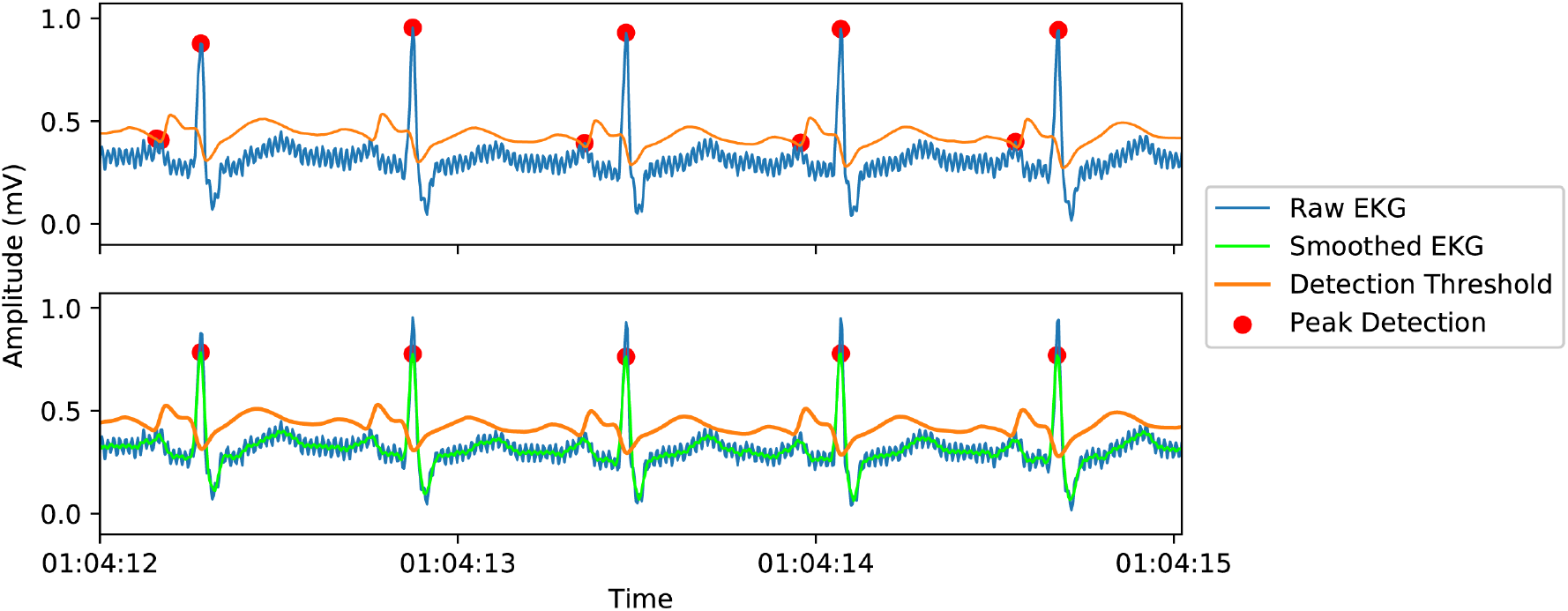
Illustration of EKG smoothing. **Top panel)** Example of a noisy EKG signal with default detection parameters, resulting in many false peak detections. **Bottom panel)** The same signal with pre-detection smoothing applied, producing marked improvement in detection accuracy. Raw EKG is shown in blue and smoothed EKG in green. Detection thresholds are shown in orange and peak detections in red.

Additionally, CardioPy can handle pre-cleaned data without error. For data with missing segments, artificially long interbeat intervals are automatically removed in the artifact rejection step, described in detail in the following section.

### 3.2 Data Inspection & Artifact Rejection

#### 3.2.1 Data Visualization

The EKG object provides self-contained methods for visualization and artifact rejection. These methods are built on the matplotlib interactive graphical user interface, which can be leveraged from a python interpreter or through a jupyter notebook.

Threshold parameters can be visualized using the *plotpeaks* method, which includes interbeat intervals with corresponding R-peak detections (**Figure 4**). Potential detection errors can be identified using the interbeat interval graph combined with matplotlib’s interactive zoom (**Figure 4, inset**) and scrollbars.

**Figure 4.**
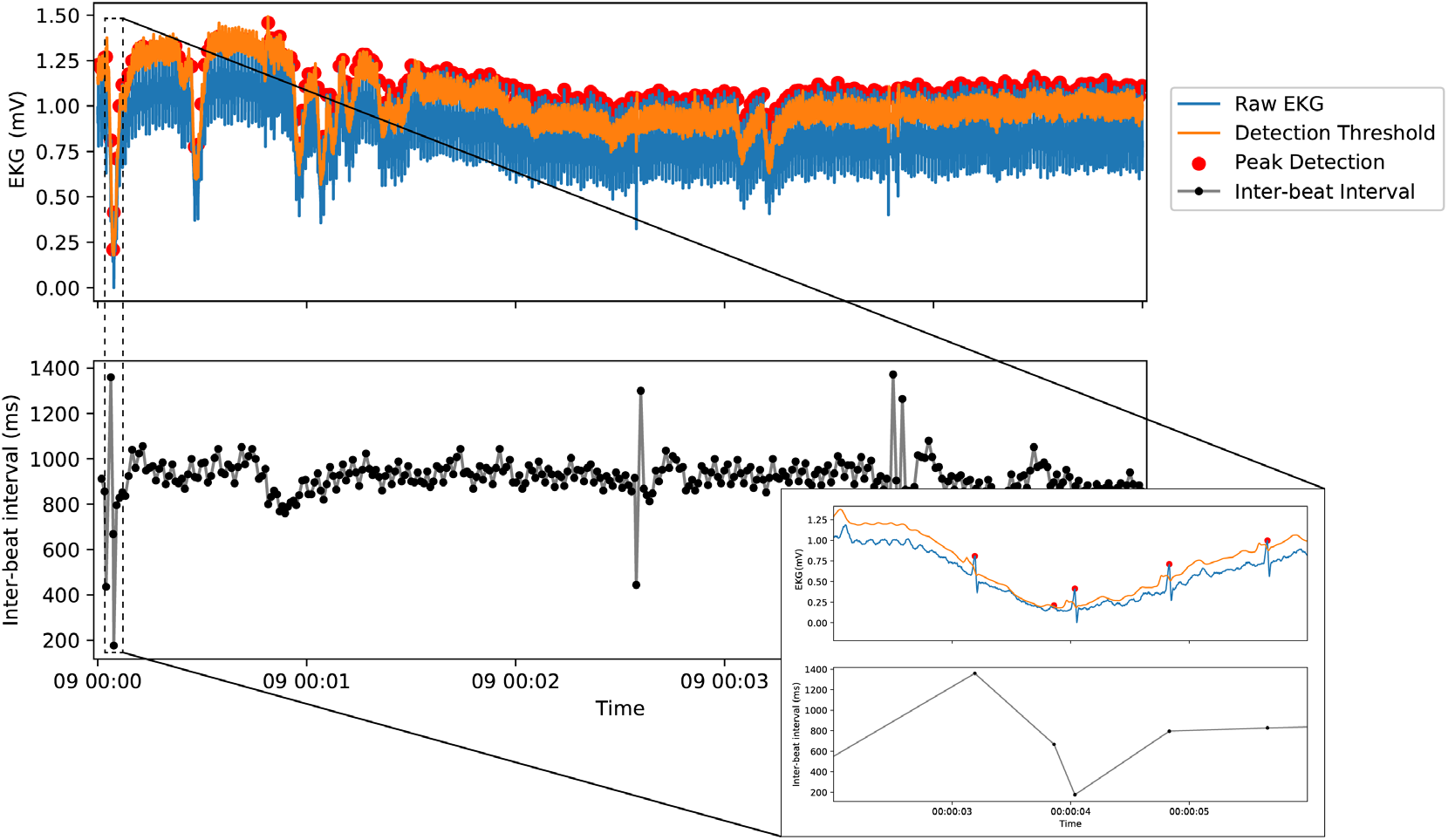
Illustration of data inspection using the *plotpeaks* method. Raw EKG signal, detection threshold, and peak detections across a 5-minute dataset **(Top Panel)** are displayed with a time-locked plot of interbeat interval lengths **(Bottom Panel)**. Irregularly short or long interbeat intervals can be inspected using the matplotlib gui’s built-in zoom and scroll functions, allowing for easy identification of missed or false R-peak detections **(Inset)**, as well as ectopic or abnormal beats.

#### 3.2.2 Artifact Rejection

False detections can be removed and missed peaks added by calling the *rm_peak* and *add_peak* methods, respectively, and specifying the time of detection error. To simplify time estimation, the exact millisecond value under the cursor is visible in either the lower left or right corner of the plotting window, depending on the matplotlib backend in use. All cleaning is tracked in artifact logs, stored as attributes of the EKG object (named rpeak_artifacts, rpeaks_added, and ibi_artifacts for false peaks, missed peaks, and removed interbeat intervals, respectively). If a true peak is mistakenly removed or a new peak added in a false position, these actions can be reversed with the *undo_rm_peak* and *undo_add_peak* methods, respectively.

#### 3.2.3 Missing Data

Missing data will result in falsely exaggerated interbeat intervals. To prevent potential dilution of true variability by interpolating missing data, we chose to handle missing data as NaN artifact. The *rm_ibi* method will automatically remove any interbeat intervals longer than a specified threshold, defaulted at 3000 milliseconds. In the case of ectopic or abnormal beats that affect multiple intervals, individual interbeat intervals can be manually specified for removal. Of note, any false or missed peaks must be corrected prior to interbeat interval artifact handling, since the former methods re-calculate interval values with each new call. Once all artifacts are removed, the resulting NN tachogram (indicating <u>N</u>ormal interbeat intervals) is used for HRV statistics.

### 3.3 HRV Analysis

Standard HRV analyses **(Table 2)** are conducted with the *hrv_stats* method (**Supplementary Code Snippet 6**). For frequency calculations, CardioPy applies cubic interpolation to the NN tachogram to produce an evenly sampled time series, which is then resampled to the sampling frequency of the original signal to minimize fiducial point shift [12]. The power spectrum is calculated using welch or multitaper methods (default: multitaper), with options to specify window type (default: Hamming) or resolution bandwidth (default: 0.008 Hz), respectively [13]. Spectra are analyzed according to standard HRV frequency bands (upper bounds inclusive) and visualized using the *plotPS* method.

### 3.4 Data Export

CardioPy EKG objects contain built-in methods for the export of cleaned data, calculations, and summary statistics. Using the *export_RR* method, peak detections and interbeat interval tachograms (cleaned or uncleaned) can be exported into plain text files compatible with major HRV analysis programs. If the RR intervals have been cleaned, this method also exports artifact logs for analytic reproducibility. The *export_RR* method can be called before, after, or without statistical analysis; accordingly, CardioPy can be used as an R-peak detector and cleaning module upstream of other software packages.

HRV analyses can also be exported into reports containing all calculated statistics for each file using the *to_report* method. The *to_spreadsheet* method writes statistics for each analyzed file to a new row of a master excel spreadsheet for group analyses.

### 3.5 CardioPy Validation & Application Example: HRV across wakefulness and sleep

#### 3.5.1 CardioPy Validation

To test the validity of the CardioPy package, we compared our analysis against Kubios HRV software [5]; Kubios is the most widely used HRV software in the scientific community, with over 1200 citations to date. For validation and the subsequent HRV example, data was collected from 5 healthy subjects using continuous 24-hour EEG with single-lead EKG at a sampling rate of 250 Hz (Natus Neuroworks, San Carlos, CA). Written informed consent was obtained from each subject for data collection and publication under protocols approved by Rockefeller University and Weill Cornell Medicine IRBs.

Sleep was scored in 30-second epochs according to standard sleep scoring criteria [14]. Each continuous period of wake, rapid-eye-movement (REM) sleep, or slow wave sleep (SWS) was divided into five-minute segments and EKG exported. For each subject, a maximum of six segments were analyzed from each brain state (mean = 4.5, SD = 1.2, contingent on availability) for a total of 66 segments. R-peaks were cleaned and interbeat intervals exported using CardioPy. HRV analysis of the NN tachogram was then conducted with CardioPy and Kubios software in parallel.

Visual inspection showed little difference in time domain results between Kubios and CardioPy. To determine the CardioPy multitaper bandwidth parameter that most closely approximated Kubios’ default power spectral estimates, we calculated the mean squared error between spectra generated by CardioPy and Kubios for each segment, using 0.0005 Hz intervals between 0.005 Hz and 0.10 Hz for CardioPy multitaper estimates (**Figure 6**). According to minimal mean squared error, 0.008 Hz was chosen as the default bandwidth parameter and applied to all subsequent analyses.

**Figure 6.**
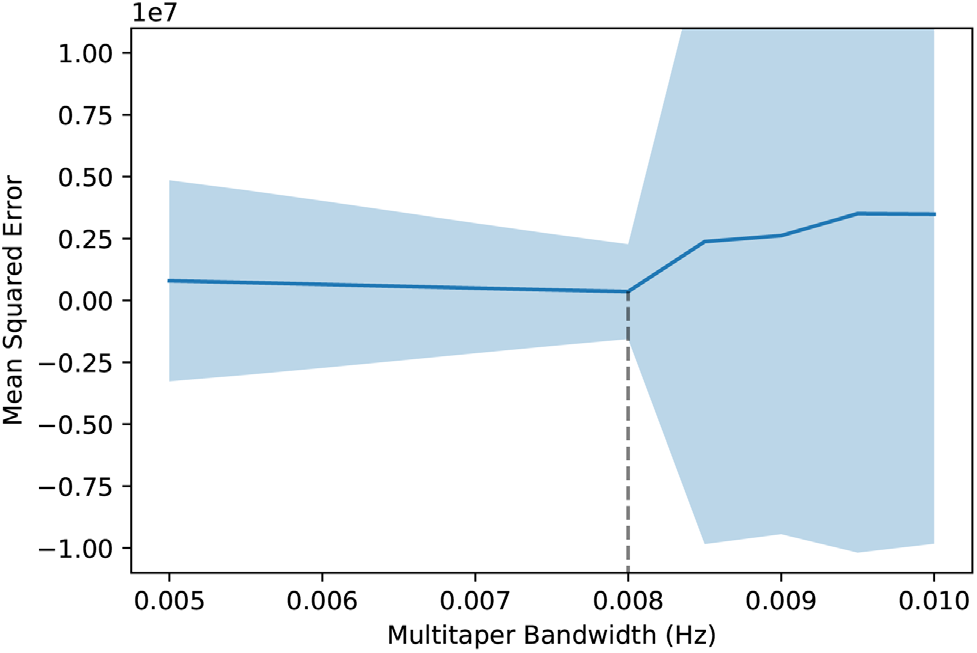
CardioPy bandwidth selection. Power spectra were calculated with CardioPy using multitaper bandwidths ranging from 0.005 to 0.010 Hz for all segments. Mean squared error was calculated against corresponding spectra generated with Kubios HRV to determine the multitaper bandwidth parameter that most closely approximated Kubios defaults. Dotted line represents minimum mean squared error at 0.008 Hz.

For each segment, we then quantitatively compared power spectra generated by each program. Default parameters were used for both CardioPy and Kubios, corresponding to multitaper estimation with 0.008 Hz bandwidth and welch estimation with 300 ms window length and 50% overlap, respectively. Across segments, the CardioPy spectra explained an average of 99.7% (SD 0.5%) of the variance present in the corresponding Kubios spectra. **Figure 7** demonstrates representative corresponding power spectra generated by CardioPy and Kubios HRV for a single EKG segment.

**Figure 7.**
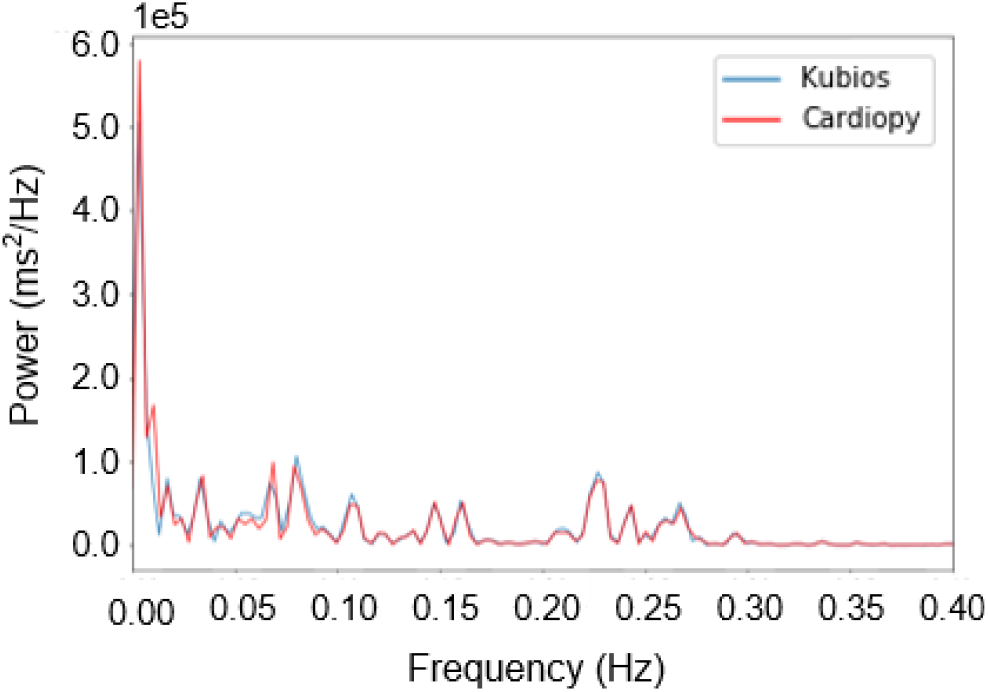
Representative power spectra generated by Kubios HRV and CardioPy from a single EKG segment. Spectra were produced using default parameters for both CardioPy (red) and Kubios HRV (blue).

#### 3.5.2 Sleep Stage Analysis

To demonstrate utility, we used CardioPy to compare HRV metrics across wakefulness, REM, and SWS. For each segment, detection parameters were optimized, data cleaned, and statistics calculated (see Supplementary Material for a step-by-step example). Optimization of detection parameters achieved a minimum detection sensitivity of 98.9% and PPV of 97.0%.

Across all segments, parameter optimization achieved average sensitivity and PPV values of 100.0% (SD 0.05%) and 99.8% (SD 0.2%), respectively (**Table 3**). Of note, since ectopic beats contain a valid R-peak in the EKG signal, they were categorized as true peaks and corresponding interbeat intervals were subsequently removed during clearning.

**Table 3.** R-Peak detection statistics. Detection parameters were optimized for each segment and results tabulated across all segments for each subject. Total values are cumulative across segments. Sensitivity and PPV values were calculated for each segment and mean value taken across segments. Standard deviations are displayed in parentheses.

| Subject | Segments<br>(Total) | True Peaks<br>(Total) | False<br>Peaks<br>(Total) | Missed<br>Peaks<br>(Total) | Sensitivity<br>(Mean) | PPV (Mean) |
| --- | --- | --- | --- | --- | --- | --- |
| 1 | 7 | 3,469 | 8 | 0 | 1.000 | 0.998 |
| 2 | 16 | 7,714 | 5 | 0 | 1.000 | 0.999 |
| 3 | 15 | 4,674 | 12 | 4 | 0.999 | 0.997 |
| 4 | 15 | 4,734 | 26 | 4 | 0.999 | 0.995 |
| 5 | 13 | 5,184 | 2 | 0 | 1.000 | 1.000 |
| Total | 66 | 25,775 | 53 | 8 | 1.000 (0.0005) | 0.998 (0.0020) |

Standard time-domain and frequency-domain metrics (outlined in **Table 2**) were compared according to brain state (Awake, REM, SWS) in a combined group analysis using Tukey’s HSD (alpha = 0.05). There were no differences between states in RMSSD, HF power or average heart rate. However, SDNN, LF power and LF/HF ratio were significantly reduced during SWS compared to awake. SDNN and LF power were also significantly reduced during SWS compared to REM sleep (**Table 4**).

**Table 4.** HRV results according to brain state for group and individual analyses. States that are significantly different according to Tukey’s HSD are grouped by superscript (*,† = *p* < 0.05; **,†† = *p* < 0.01; ***, ††† = *p* ≤ 0.001). Where only a single segment was available for analysis, no standard deviation value is listed. N/A values indicate that no segments were available for analysis.

|  | Group | Subject 1 | Subject 2 | Subject 3 | Subject 4 | Subject 5 |
| --- | --- | --- | --- | --- | --- | --- |
| **RMSSD** |  |  |  |  |  |  |
| Awake | 38 ± 25 | 10 ± 2** | 33 ± 15* | 64 ± 8 | 60 ± 27* | 32 ± 15*** |
| REM | 49 ± 29 | 24** | 17 ± 7 | 66 ± 14 | 52 ± 18 | 88 ± 20***†† |
| SWS | 40 ± 26 | N/A | 11 ± 8* | 75 ± 10 | 24 ± 8* | 50 ± 6†† |
| **SDNN** |  |  |  |  |  |  |
| Awake | 56 ± 28* | 23 ± 2*** | 57 ± 25* | 76 ± 18 | 79 ± 20** | 51 ± 23** |
| REM | 65 ± 26†† | 47*** | 40 ± 9 | 85 ± 21 | 63 ± 22† | 96 ± 8**†† |
| SWS | 36 ± 17*†† | N/A | 18 ± 8* | 58 ± 9 | 28 ± 7**† | 42 ± 4†† |
| **LF power (ms<sup>2</sup> × 10<sup>4</sup>)** |  |  |  |  |  |  |
| Awake | 37 ± 41* | 5 ± 2*** | 46 ± 52 | 49 ± 16 | 73 ± 61 | 19 ± 12*** |
| REM | 39 ± 31† | 28*** | 9 ± 6 | 56 ± 24* | 43 ± 37 | 68 ± 22***††† |
| SWS | 11 ± 8*† | N/A | 4 ± 3 | 21 ± 10* | 8 ± 4 | 10 ± 4††† |
| **HF power (ms<sup>2</sup> × 10<sup>4</sup>)** |  |  |  |  |  |  |
| Awake | 18 ± 19 | 0.8 ± 0.3*** | 12 ± 11 | 38 ± 9 | 33 ± 26 | 11 ± 12*** |
| REM | 33 ± 33 | 11*** | 3 ± 3 | 53 ± 23 | 22 ± 16 | 81 ± 29***†† |
| SWS | 26 ± 27 | N/A | 2 ± 3 | 63 ± 21 | 6 ± 3 | 32 ± 7†† |
| **LF/HF** |  |  |  |  |  |  |
| Awake | 3.5 ± 2.8** | 7.0 ± 3.1 | 4.0 ± 2.6 | 1.3 ± 0.3* | 2.2 ± 0.4 | 2.2 ± 0.7**††† |
| REM | 2.0 ± 1.2 | 2.5 | 3.3 ± 1.3 | 1.2 ± 0.7† | 2.1 ± 0.7 | 0.9 ± 0.2** |
| SWS | 1.5 ± 2.1** | N/A | 4.5 ± 4.2 | 0.3 ± 0.1*† | 1.5 ± 0.6 | 0.3 ± 0.1††† |
| **HR avg (bpm)** |  |  |  |  |  |  |
| Awake | 73 ± 9 | 80 ± 4 | 76 ± 5** | 65 ± 2* | 63 ± 4 | 82 ± 8**† |
| REM | 69 ± 9 | 82 | 81 ± 4 | 63 ± 2 | 63 ± 2 | 66 ± 1** |
| SWS | 72 ± 11 | N/A | 89 ± 3** | 60 ± 3* | 69 ± 5 | 72 ± 0† |

To examine the robustness of these findings, we then compared between-state differences at the individual subject level. Of note, group trends did not always hold true; no single metric was found to consistently vary in all subjects. However, RMSSD and SDNN were both significantly reduced during SWS compared to wake in 3 of 4 subjects where SWS was present. LF power was less consistent, showing a significant increase from wake to REM in 2 of 5 subjects, and reduction from REM to SWS in 2 of 4 subjects. Between-state differences in HF power and LF/HF ratio were found in only 2 subjects, while average heart rate significantly varied between wake and SWS in 3 of 4 subjects, although in differing directions (**Table 4**).

## 4. DISCUSSION

Here we present a novel open-source package for the analysis of single-lead EKG data. Our goal was to reduce the barrier to entry for HRV research using clinical EKG signals by introducing a simple, free, OS agnostic and easy-to-implement analysis toolkit. CardioPy requires minimal programming experience and can be run interactively through Jupyter Notebook, leveraging the built-in user-friendly functionalities of both jupyter and matplotlib, while also facilitating a high level of scientific reproducibility. We provide thorough documentation of the functionalities and use of the CardioPy toolkit, as well as demonstrate analytic validity through comparison against Kubios HRV and example analysis of HRV in different brain states.

### 4.1 Performance & Validation

In HRV analyses, minor artifacts in R-peak detection can produce significant skewing of the NN tachogram and corresponding HRV statistics. For this reason, and in accordance with the recommendations of the European Task Force (1996), we made the strategic decision to exclude automatic artifact detection from the CardioPy program. While established detectors have reported excellent accuracy with automatic artifact detection algorithms, in our experience many automatic detection methods had the conservative effect of washing out a portion of true variability in our data. Instead, we implemented a highly flexible detection algorithm in combination with manual data inspection and artifact rejection.

From our example analyses, we show that adaptation of the detection parameters to varying signal-to-noise ratios produced an average detection sensitivity of 100.0% and PPV of 99.8% (**Table 3**). The few remaining artifacts were removed with CardioPy’s built-in artifact removal methods. A potential disadvantage of this method is the time required for manual parameter adjustment and artifact removal. We found parameter optimization to be the most time-intensive step of this approach, although for no dataset did parameter optimization require more than 2-3 minutes. In future improvements on the CardioPy algorithm we plan to streamline this step with automatic parameter adjustments based on the signal-to-noise characteristics of each dataset. Additionally, although manual artifact rejection is slower than automated options, manual rejection requires a high degree of data inspection and ensures the overall quality of the resultant NN tachogram, which is critical for accurate and reproducible results.

We further validated CardioPy’s calculations against the Kubios HRV software. CardioPy spectra closely approximated those produced by Kubios, explaining approximately 99.7% (SD 0.5%) variance of the Kubios spectra. Final time and frequency domain HRV statistics similarly showed minimal variation, demonstrating the statistical validity of the CardioPy analysis pipeline.

### 4.2 Example Application: HRV in Wakefulness and Sleep

In our HRV statistics calculated across waking, REM, and SWS states in 5 healthy subjects, we show results consistent with reported sleep-related changes in autonomic tone from waking into different stages of sleep [15,16]. Specifically, we found that LF power and SDNN were significantly reduced during SWS sleep compared to awake and REM sleep states (**Table 4**). Similar to what has been reported in other studies of healthy individuals [17,18], calculated LF/HF ratios were significantly lower during SWS sleep compared to awake states (**Table 4**), likely reflecting an increase in parasympathetic tone with increasing sleep depth. This metric is fairly robust against variations in power spectral calculation methods and has shown clinical utility across a variety of conditions [19–21].

Notably, we also underscore the individual variability of HRV analyses, finding that group-level differences were largely driven by highly significant changes in a minority of subjects (**Table 4**). This result highlights the caution that must be taken with HRV interpretation; we suggest future studies on intra-individual HRV analyses account for potentially mediating factors that may affect autonomic regulation, such as time of night [22].

### 4.2 Limitations

A significant current limitation of the CardioPy toolkit is the lack of a graphical user interface. Although CardioPy utilizes matlplotlib’s interactive functionality, analyses must be run through the command line, requiring the minimal programming capabilities of setting up an Anaconda distribution and operating within the jupyter interface. To mitigate this limitation, we have provided a template jupyter notebook to produce a plug-and-play implementation. In future versions of CardioPy, we hope to further address this limitation by developing a more approachable graphical user interface.

Secondly, CardioPy is optimized for short-term recordings and may not be appropriate for longer datasets. The CardioPy pipeline relies heavily on Pandas data structures, which are efficient for five-minute datasets but produce significant computational overhead and may slow with increased file sizes. CardioPy also does not include calculations that require long-term recordings, such as ULF power and detrended fluctuation analysis. As short-term recordings are currently the most common form of HRV data in published studies [10], this limitation is unlikely to significantly affect usability.

### 4.3 Future Directions

In future versions of CardioPy, we hope to implement a graphical user interface, as well as support for additional commonly used data formats. Although the decision against implementing an automatic artifact rejection algorithm increases the time required for data cleaning, we feel it has the potential to improve the accessibility and quality of clinical HRV research by (1) being computationally simple and easy to understand, and (2) requiring a degree of data inspection for each analysis. Furthermore, we intend to streamline the CardioPy algorithm by implementing automatic detection parameter adjustment, which will expedite this preprocessing step.

CardioPy provides the contribution of a freely available R-peak detection tool that is not restricted to the Windows operating system and is not reliant on the proprietary MATLAB environment. It is an open-source program, and we encourage collaboration and improvements from the larger community of researchers and programmers. We hope that CardioPy provides a useful addition to the growing set of excellent resources that facilitate meaningful clinical research.

## Supporting information

Supplemental Code Snippets

## Acknowledgements

The authors thank Dr. Jonathan Victor for advice regarding data analysis.

## REFERENCES

[1] F. Shaffer, R. McCraty, C.L. Zerr, A healthy heart is not a metronome: an integrative review of the heart’s anatomy and heart rate variability, Front. Psychol. 5 (2014) 1–19. https://doi.org/10.3389/fpsyg.2014.01040.

[2] D.J. Plews, P.B. Laursen, J. Stanley, A.E. Kilding, M. Buchheit, Training Adaptation and Heart Rate Variability in Elite Endurance Athletes: Opening the Door to Effective Monitoring, Sport. Med. 43 (2013) 773–781. https://doi.org/10.1007/s40279-013-0071-8.

[3] H.G. Kim, E.J. Cheon, D.S. Bai, Y.H. Lee, B.H. Koo, Stress and heart rate variability: A meta-analysis and review of the literature, Psychiatry Investig. 15 (2018) 235–245. https://doi.org/10.30773/pi.2017.08.17.

[4] R.P. Villareal, B.C. Liu, A. Massumi, Heart rate variability and cardiovascular mortality, Curr. Atheroscler. Rep. 4 (2002) 120–127. https://doi.org/10.1007/s11883-002-0035-1.

[5] M.P. Tarvainen, J.P. Niskanen, J.A. Lipponen, P.O. Ranta-aho, P.A. Karjalainen, Kubios HRV - Heart rate variability analysis software, Comput. Methods Programs Biomed. 113 (2014) 210–220. https://doi.org/10.1016/j.cmpb.2013.07.024.

[6] L. Rodríguez-Liñares, M.J. Lado, X.A. Vila, A.J. Méndez, P. Cuesta, gHRV: Heart rate variability analysis made easy, Comput. Methods Programs Biomed. 116 (2014) 26–38. https://doi.org/10.1016/j.cmpb.2014.04.007.

[7] V. Pichot, F. Roche, S. Celle, J. Barthélémy, F. Chouchou, HRVanalysis: A Free Software for Analyzing Cardiac Autonomic Activity, Front. Physiol. 7 (2016) 1–15. https://doi.org/10.3389/fphys.2016.00557.

[8] T. Kaufmann, S. Sütterlin, S.M. Schulz, C. Vögele, ARTiiFACT: a tool for heart rate artifact processing and heart rate variability analysis, Behav. Res. Methods. 43 (2011) 1161–1170. https://doi.org/10.3758/s13428-011-0107-7.

[9] J.E. Muñoz, E.R. Gouveia, M.S. Cameirão, S.B. i Badia, PhysioLab - a multivariate physiological computing toolbox for ECG, EMG and EDA signals: a case of study of cardiorespiratory fitness assessment in the elderly population, Multimed. Tools Appl. 77 (2018) 11521–11546. https://doi.org/10.1007/s11042-017-5069-z.

[10] F. Shaffer, J.P. Ginsberg, An Overview of Heart Rate Variability Metrics and Norms, Front. Public Heal. 5 (2017) 1–17. https://doi.org/10.3389/fpubh.2017.00258.

[11] T.F. of T.E.S. of C. and T.N.S. of P. and American, Electrophysiology, Heart rate variability: Standards of measurement, physiological interpretation, and clinical use Task, Eur. Heart J. 17 (1996) 354–381. http://search.ebscohost.com/login.aspx?direct=true&db=ufh&AN=1888969&site=ehost-live.

[12] G.D. Clifford, L. Tarassenko, Quantifying errors in spectral estimates of HRV due to beat replacement and resampling, IEEE Trans. Biomed. Eng. 52 (2005) 630– 638. https://doi.org/10.1109/TBME.2005.844028.

[13] A. Gramfort, M. Luessi, E. Larson, D.A. Engemann, D. Strohmeier, C. Brodbeck, R. Goj, M. Jas, T. Brooks, L. Parkkonen, M. H??m??l??inen, MEG and EEG data analysis with MNE-Python, Front. Neurosci. 7 (2013) 1–13. https://doi.org/10.3389/fnins.2013.00267.

[14] Q.S. Iber C, Ancoli-Israel S, Chessonn A, The AASM Manual for the scoring of sleep and associated events: rules, terminology and technical specifications, 1st ed., American Academy of Sleep Medicine, Westchester, IL, 2007.

[15] E. Tobaldini, L. Nobili, S. Strada, K.R. Casali, A. Braghiroli, N. Montano, Heart rate variability in normal and pathological sleep, Front. Physiol. 4 (2013) 1–11. https://doi.org/10.3389/fphys.2013.00294.

[16] M. de Zambotti, J. Trinder, A. Silvani, I.M. Colrain, F.C. Baker, Dynamic coupling between the central and autonomic nervous systems during sleep: A review, Neurosci. Biobehav. Rev. 90 (2018) 84–103. https://doi.org/10.1016/j.neubiorev.2018.03.027.

[17] D. Herzig, P. Eser, X. Omlin, R. Riener, M. Wilhelm, P. Achermann, Reproducibility of heart rate variability is parameter and sleep stage dependent, Front. Physiol. 8 (2018) 1–10. https://doi.org/10.3389/fphys.2017.01100.

[18] P. Boudreau, W.-H. Yeh, G.A. Dumont, D.B. Boivin, Circadian Variation of Heart Rate Variability Across Sleep Stages, Sleep. 36 (2013) 1919–1928. https://doi.org/10.5665/sleep.3230.

[19] H. Cohen, J. Benjamin, A.B. Geva, M.A. Matar, Z. Kaplan, M. Kotler, Autonomic dysregulation in panic disorder and in post-traumatic stress disorder: Application of power spectrum analysis of heart rate variability at rest and in response to recollection of trauma or panic attacks, Psychiatry Res. 96 (2000) 1–13. https://doi.org/10.1016/S0165-1781(00)00195-5.

[20] H. Cohen, L. Neumann, M. Shore, M. Amir, Y. Cassuto, D. Buskila, Autonomic dysfunction in patients with fibromyalgia: Application of power spectral analysis of heart rate variability, Semin. Arthritis Rheum. 29 (2000) 217–227. https://doi.org/10.1016/S0049-0172(00)80010-4.

[21] M. Patel, S.K.L. Lal, D. Kavanagh, P. Rossiter, Applying neural network analysis on heart rate variability data to assess driver fatigue, Expert Syst. Appl. 38 (2011) 7235–7242. https://doi.org/10.1016/j.eswa.2010.12.028.

[22] A. Kontos, M. Baumert, K. Lushington, D. Kennedy, M. Kohler, D. Cicua-navarro, Y. Pamula, J. Martin, The Inconsistent Nature of Heart Rate Variability During Sleep in Normal Children and Adolescents, 7 (2020) 1–11. https://doi.org/10.3389/fcvm.2020.00019.

