## Supplemental Code Snippets for "CardioPy: An open-source heart rate variability toolkit for single-lead EKG"

**Supplementary Material**

Step-by-step code snippets presented below represent the full HRV analysis of a single 5-minute segment, utilizing interactive plotting in a jupyter notebook environment. If working inside a jupyter notebook, interactive plots should be enabled using one of two matplotlib magic commands. For traditional pop-out plots, the first command within the jupyter notebook should be *%matplotlib qt*, whereas for embedded plots the notebook should begin with *%matplotlib notebook*.

<u>Step 1: Load data and detect peaks</u>. CardioPy automatically conducted R-peak detection upon data loading and initialization of the EKG object. Any adjustments to detection threshold parameters after viewing default detections were implemented by re-initializing the EKG object with new *upshift* or *mw_size* parameters. The example 5-minute epoch was initialized with a 2% *upshift* and 55 millisecond *mw_size* (**Code Snippet 1**).

<u>Step 2. Visualize and clean peak detections</u>. Detections were visualized with the *plotpeaks* method (**Code Snippet 2**). Close inspection revealed 1 missed peak and 1 false peak, which were corrected with the *add_peak* and *rm_peak* methods, respectively (**Code Snippets 3 & 4**). The *plotpeaks* method was called iteratively to inspect initial detections and parameter adjustments, as well as R-peak cleaning accuracy. Any irregular interbeat intervals produced by ectopic or otherwise abnormal beats were removed with the *rm_ibi* method. Once cleaning was complete, processed R-peaks and NN intervals were exported with the *export_RR* method (**Code Snippet 5**).

Step 3. Calculate and export HRV statistics. HRV statistics were calculated with a call to the *hrv_stats* method and power spectrum visualized with the *plotPS* method (**Code Snippet 6**). Data was exported to a single report as well as a cumulative spreadsheet (**Code Snippet 7**).

Set interactive plotting and import the EKG class

```
%matplotlib notebook
from cardiopy.ekg import EKG
```

Specify the data file, path to file, and directory to save output

```
fname = 'HCXXX_2001-01-01_sws_cycle0_epoch0_000000.csv'
fpath, savedir = '.', '..'
```

Create the EKG object 'e' and detect peaks

```
e = EKG(fname, fpath, upshift=2, mw_size=55)
```

```
EKG successfully imported.
Calculating moving average with 55 ms window and a 2% upshift...
Detecting R peaks...
R peak detection complete
R-R intervals calculated
```

**Code Snippet 1 | Data loading and peak detection**

```
e.plotpeaks()
```

```
<IPython.core.display.Javascript object>
```

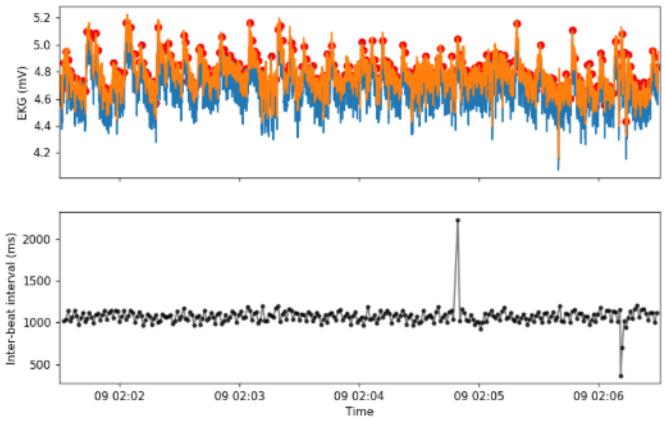


**Code Snippet 2 | Visualization of peak detections (top panel) and interbeat intervals (bottom panel)**

```
e.plotpeaks()
```

```
<IPython.core.display.Javascript object>
```

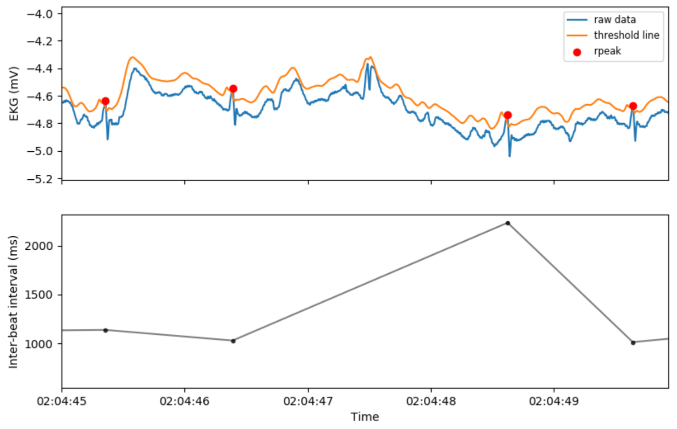


```
e.add_peak('02:04:48')
```

```
Millisecond range of missed peak [min:max]: 300:350
New peak added.
ibi values recalculated.
```

**Code Snippet 3 | Addition of missed peaks**

```
e.plotpeaks()
```

```
<IPython.core.display.Javascript object>
```

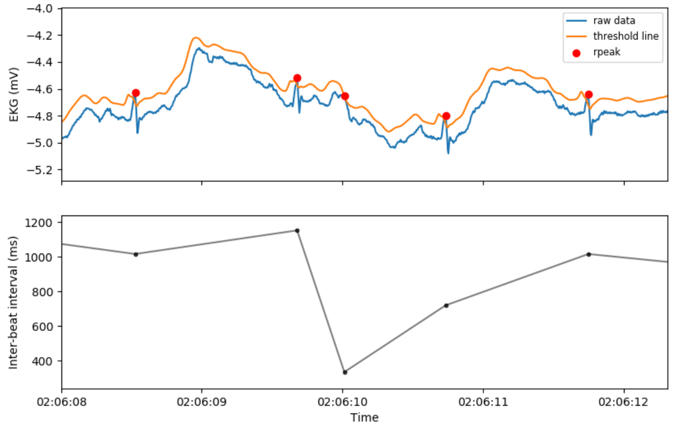


```
e.rm_peak('02:06:11')
```

```
id        time                          ibi_ms
1         2014-01-09 02:06:11.036000    364.0
2         2014-01-09 02:06:11.732000    696.0
Rpeaks to remove [list ids or None]: 1


R peak at  2014-01-09 02:06:11.036000  successfully removed.
ibi values recalculated.
```

**Code Snippet 4 | Removal of false peaks**

```
e.export_RR(savedir)
```

```
Files will be saved to ..
R peak artifacts exported.
R peak additions exported.
R peaks exported.
IBI artifacts exported.
rr intervals exported.
nn intervals exported.
Done.
```

**Code Snippet 5 | Cleaned data export.**

```
e.hrv_stats()
```

```
Calculating time domain statistics...
Time domain stats stored in obj.time_stats

Interpolating and resampling tachogram...
Calculating power spectrum...
Calculating frequency domain measures...
Frequency measures stored in obj.freq_stats

Done.
```

```
fig = e.plotPS(save = False)
```

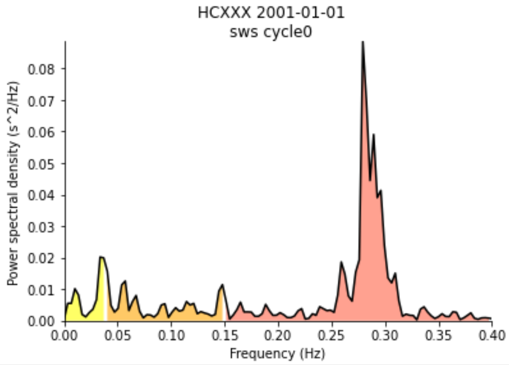


**Code Snippet 6 | HRV statistics calculation & power spectrum visualization.** Spectrum is colored by frequency band (Yellow: VLF, Orange: LF, Red: HF)

```
e.to_report(savedir)
```

```
Files will be saved to ..
```

```
e.to_spreadsheet('HRV_Stats_Spreadsheet.csv', savedir)
```

```
Data added to HRV_Stats_Spreadsheet.csv
```

**Code Snippet 7 | HRV report and spreadsheet export**
